# Event-related causality in Stereo-EEG discriminates syntactic processing of noun phrases and verb phrases

**DOI:** 10.1101/2022.02.25.481939

**Authors:** Andrea Cometa, Piergiorgio D’Orio, Martina Revay, Franco Bottoni, Claudia Repetto, Giorgio Lo Russo, Stefano F. Cappa, Andrea Moro, Silvestro Micera, Fiorenzo Artoni

## Abstract

Syntax involves complex neurobiological mechanisms, which are difficult to disentangle for multiple reasons. Using a protocol able to separate syntactic information from sound information we investigated the neural causal connections evoked by the processing of homophonous phrases, either verb phrases (VP) or noun phrases (NP). We used event-related causality (ERC) from stereo-electroencephalographic (SEEG) recordings in 10 epileptic patients in multiple cortical areas, including language areas and their homologous in the non-dominant hemisphere. We identified the different networks involved in the processing of these syntactic operations (faster in the dominant hemisphere) showing that VPs engage a wider cortical network. We also present a proof-of-concept for the decoding of the syntactic category of a perceived phrase based on causality measures. Our findings help unravel the neural correlates of syntactic elaboration and show how a decoding based on multiple cortical areas could contribute to the development of speech prostheses for speech impairment mitigation.

## Introduction

Traditionally, language is analyzed in relation to four main components: the acoustic level, that is the physical medium humans naturally exploit to convey information and its articulatory–phonatory counterpart; the lexicon, which is the repertoire of words expressing predicative contents and logical instructions; syntax, the set of principles to assemble larger units (phrases) from lexical items, in a recursive potentially infinite way; semantics, an interpretative component which captures the truth value conditions for each syntactic structure. However, since the acoustic and syntactic information are crucially intertwined (Ding et al., 2015), even during inner speech (Kayne, 2019; Magrassi et al., 2015), isolating syntax at the electrophysiological level appears to be an insurmountable empirical task. This is reflected in the difficulty of developing specific syntax-related tasks for experimental studies of language neurobiology and it is responsible for the relatively limited knowledge of syntax-related processing in the brain. Understanding the neural correlates of even the most basic syntactic operations, such as merging an article with a noun (N) yielding a Noun Phrase (NP) or a pronoun with a verb (V) yielding a Verb Phrase (VP) remains a crucial challenge for brain and language research (Grodzinsky & Friederici, 2006).

In a recent study (Artoni et al., 2020), we designed and used a novel protocol aimed at isolating syntactic information from the acoustic associated information by exploiting pairs of sentences containing homophonous strings (same acoustic information but completely different syntactic content). Specifically, each pair of stimuli contained the same acoustic copy of two homophonous words, which could be interpreted either as a Noun Phrase (NP) or a Verb Phrase (VP) **Figure 1A**). This approach was used to factor out any phonological and prosodical clue in a complete way, even at the subliminal level. We used this protocol while recording the related cortical activation using stereo-electroencephalography (SEEG), an invasive recording technique with unparalleled signal-to-noise ratio and recording band-width (He et al., 2019; Lachaux et al., 2003).

**Figure 1.**
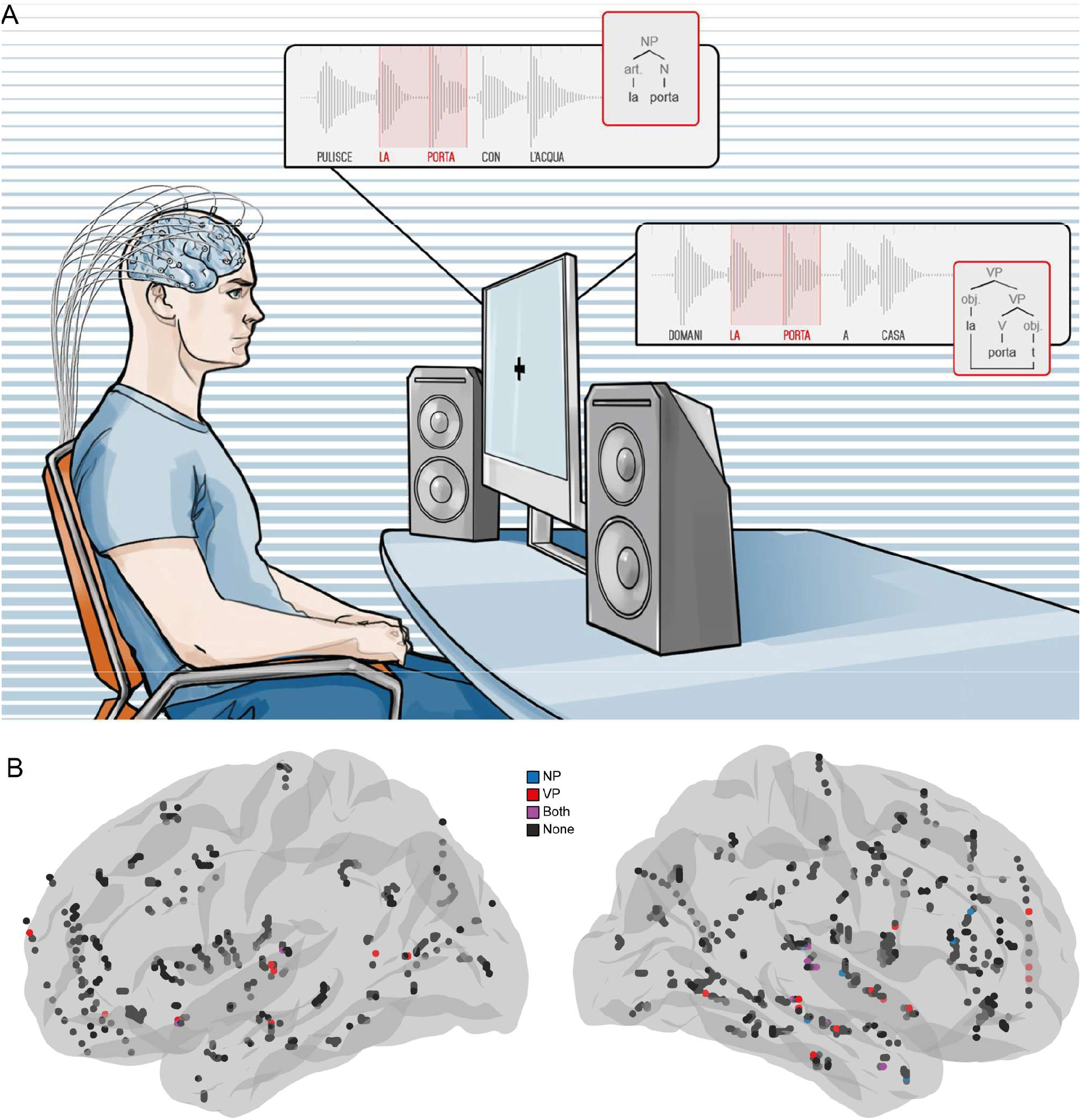
Experimental set-up. **(A)** Example of a set of homophonous sequences (i.e., strings of words with the same sound but different syntactic structure) used in the experiment. For example, in PULISCE LA PORTA CON L’ACQUA (s/he cleans the door with water), the phonemic sequence [la’pɔrta] (written here as: la porta) is a Noun Phrase (NP), while in DOMANI LA PORTA A CASA (tomorrow s/he brings her home), the same sequence is a Verb Phrase (VP). From (Artoni et al., 2020). **(B)** Mini-region-of-interests (merged across all subjects) in the dominant (left) and non-dominant (right) hemispheres. Contacts involved in the NP-related network are highlighted in blue, those involved in the VP processing network are highlighted in red, and those participating in both networks are coloured in purple.

Here, we exploited the same dataset to investigate the amplitude, the direction, and the specific frequencies of the interactions taking place between brain structures, that is the collection of causal links elicited by different functional situations known as effective connectivity (Penny et al., 2004). Given the utmost importance of timing, here we analyze the directed connectivity patterns elicited by a stimulus, i.e., the event-related causality (ERC). We investigated the dynamical evolution of the causal integration in response to a specific part of the time-varying stimuli (sentences) - the response window (RW) - either the NP or the VP. To reach this aim and to characterize and define the different networks involved in the processing of the syntactic operations yielding a Noun Phrase or a Verb Phrase we used a recently validated pipeline of ours for the evaluation of ERC in a set RW (Cometa et al., 2021).

We also present a proof-of-concept for the decoding of the syntactic category of a perceived sentence based on causality measures which could contribute to the future development of speech prostheses for speech impairment mitigation.

## Results

### NPs and VPs elicit two unique networks

The neural networks elicited by the processing of NPs and VPs were investigated with SEEG. The data were recorded from 10 Italian-native speaker patients with no language disorders who underwent surgical operation for drug-resistant epilepsy. NPs and VPs were encoded in the same acoustic stimulus and could be differentiated only by their syntactic context (some Italian homophonous phrases, such as *la porta* /la ‘pɔrta/ – that can be interpreted either as a noun phrase – “the door” – or a verb phrase – “[s/he] brings her”). After pre-processing, close recording contacts were arranged in groups called mini-regions of interest (mini-ROIs), each represented by a prototypical contact. The grouping resulted in a total of 396 mini-ROIs in the left – or dominant – hemisphere (DH) and 577 mini-ROIs in the right – or non-dominant – hemisphere (NDH) (**Figure 1B**). To identify the networks involved in both NPs and VPs processing (i.e., the group of mini-ROIs bounded together by causal relations), we used Partial Directed Coherence (PDC) (Baccalá & Sameshima, 2001) and a recently developed pipeline to determine the significance of ERC elicited by an RW (Cometa et al., 2021).

We restricted the analysis to connections identified within the ultra-high gamma frequency band (150 to 300 Hz) (Artoni et al., 2020). The pipeline discovered 13 significant connections for the NP case (2 in the DH and 11 in the NDH) and 20 connections for the VP condition (6 in the DH, 13 in the NDH, and 1 from the right temporal lobe to the left temporal lobe). We observed 4 connections active for both phrases in the NDH. Of these shared connections 3 were intra-temporal (**Figure 2A**). All the significant connections are shown in **Table S1**.

**Figure 2.**
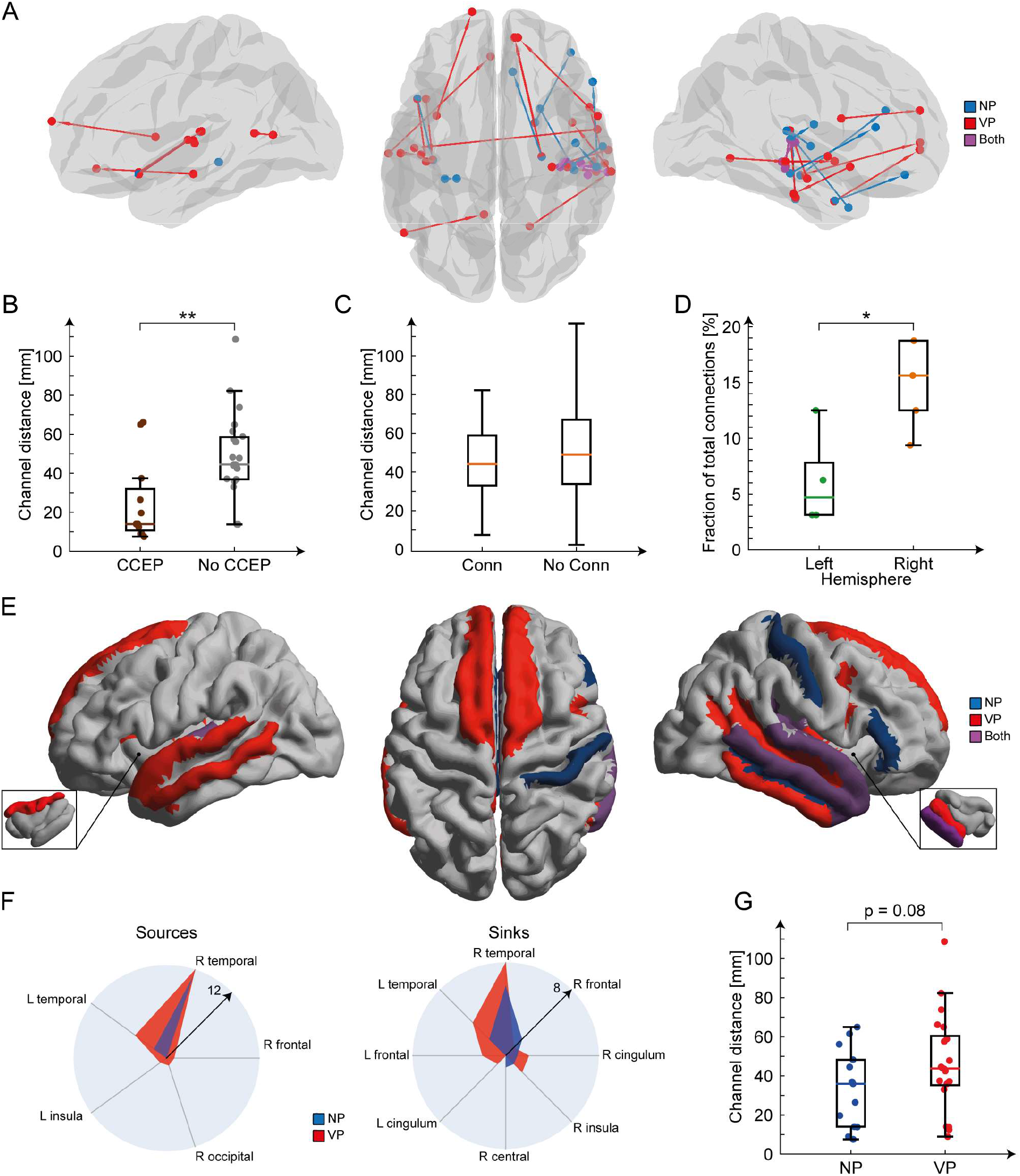
VPs engage a wider and more complex network. **(A)** Lateral and dorsal views of the identified directed connections. Nodes and edges are highlighted in blue for the Noun Phrase (NP) processing network, in red for the Verb Phrase related network, or purple if shared by both processing systems. **(B)** Box plots of the distances between the contacts involved in a significant connection and with a relevant cortico-cortical evoked potential (CCEP) and between those not showing CCEPs. **(C)** Box plots of the distances between pairs of explored contacts, whether a significant connection exists between them (Conn) or not (No Conn). **(D)** Box plots of the number of connections in subjects with electrodes in the non-dominant (right) hemisphere and in those in which only the dominant (left) was probed. The vertical axis is normalized by the total number of significant directed connections identified across all subjects. **(E)** Lateral and dorsal views of the active brain zones during NPs (blue) processing, VPs (red) processing, or both (purple). An active brain zone is a cortical area containing one or more recording contacts that act as sources or sinks for a certain directed connection. The zoom-in pictures show the left and right insula. **(F)** Radar plots of the number of sources (left) and sinks (right) in each cerebral lobe, for the two conditions NP (blue) and VP (red). **(G)** Box plots of the distances between contacts involved in a significant connection during NP and VP processing.

We compared the estimated connections with the recorded cortico-cortical evoked potentials (CCEPs) (Matsumoto & Kunieda, 2019), which are an indicator of the presence of a direct cortico-cortical or cortico-subcortico-cortical anatomical pathway (Matsumoto et al., 2004). Out of all the pairs of channels with a significant connection, only 11 exhibited a CCEP. The contacts involved in a significant connection and with a relevant CCEP were placed closer together than those not showing CCEPs (Mann-Whitney U_22,11_ = 53, p < 0.005) (**Figure 2B**).

Significant connections may be biased by clusters of closely placed contacts. Thus, to factor out a possible effect of this spatial sampling bias, we compared the distribution of the distances between pairs of contacts showing significant causal connections with the distribution of the distances between all channels (**Figure 2C**). We did not detect any difference between the two distributions (Mann-Whitney U_29,47987_ = 590819, p = 0.16).

Finally, more significant connections in both NPs and VPs were found in subjects with electrodes placed in the NDH, in contrast to those with the DH explored (Mann-Whitney U_4,5_ = 18.5, p < 0.05, **Figure 2D**). This difference was still present even when normalizing the number of significant directed connections by the total amount of the possible connections for each subject (Mann-Whitney U_4,5_ = 18, p < 0.05). Only one subject had both hemispheres explored and showed an inter-hemispheric connection (VP, from the right temporal lobe to the left one).

### VPs engage a wider network than NPs

The recording contacts participating in the NP-related network or the VP-related network were not spread across the entire cortical surface but rather clustered in specific brain zones – i.e. the anatomical parcellation of cortical gyri and sulci according to the Destrieux atlas (Destrieux et al., 2010). In total, 64 brain zones were probed in the DH and 88 in the NDH. Out of 152 cortical areas, 11 were involved in the processing of both homophonous phrases (2 in the DH and 9 in the NDH), 12 participated in the processing of the VPs alone (6 in the DH and 6 in the NDH) and 6 responded exclusively to NPs (1 in the DH and 5 in the NDH) (**Figure 2E**).

The connectivity estimated by the PDC is a directed causal information flow from one recording contact called source to another denoted sink. For NPs, all the sources were located bilaterally in the temporal lobes (2 in the DH and 11 in the NDH). For VPs, the temporal lobes contained 17 sources (5 in the DH and 12 in the NDH). The other 3 VPs sources were situated in the right occipital lobe, right frontal lobe, and left insula (**Figure 2F, left**). Most sinks, for both NPs and VPs, were in the two temporal lobes (DH: 2 for NPs and 4 for VPs; NDH: 6 for NPs and 8 for VPs). Other sinks were in the right insula (1 for NPs, 2 for VPs), in the right frontal lobe (2 for NPs, 1 for VPs), right central lobe (1 for NPs), right cingulum (1 for NPs, 2 for VPs), left frontal lobe (2 for VPs), and left cingulum (1 for VPs) (**Figure 2F, right**). The lists of the cortical areas containing sources and sinks for a given connection are shown in **Table S2** and **Table S3**.

Overall, VPs elicited more sources or sinks than NPs, engaged a higher number of different cortical areas in both hemispheres, with almost no brain-zone being more active for NPs.

The results show that VPs extended the processing network beyond the temporal lobes.

Recording contacts that participated in VPs processing seemed to be located further than those involved in NPs processing (Mann-Whitney U_13,20_ = 93, p = 0.08, **Figure 2G**), even if not reaching the statistical significance level *α* = 0.05.

### Syntax processing is faster in the DH

We then looked at the speed of response, or processing time, in the DH and NDH. The latencies of the peaks in the temporal evolutions of the time-varying significant causalities were thus compared among hemispheres. We considered only the highest peak, for each time series, occurring during the homophonous part of the stimuli (**Figure 3A**). These peaks arose earlier in the DH (Mann-Whitney U_8,24_ = 54.5, p < 0.05), for both NPs and VPs **(Figure 3B**).

**Figure 3.**
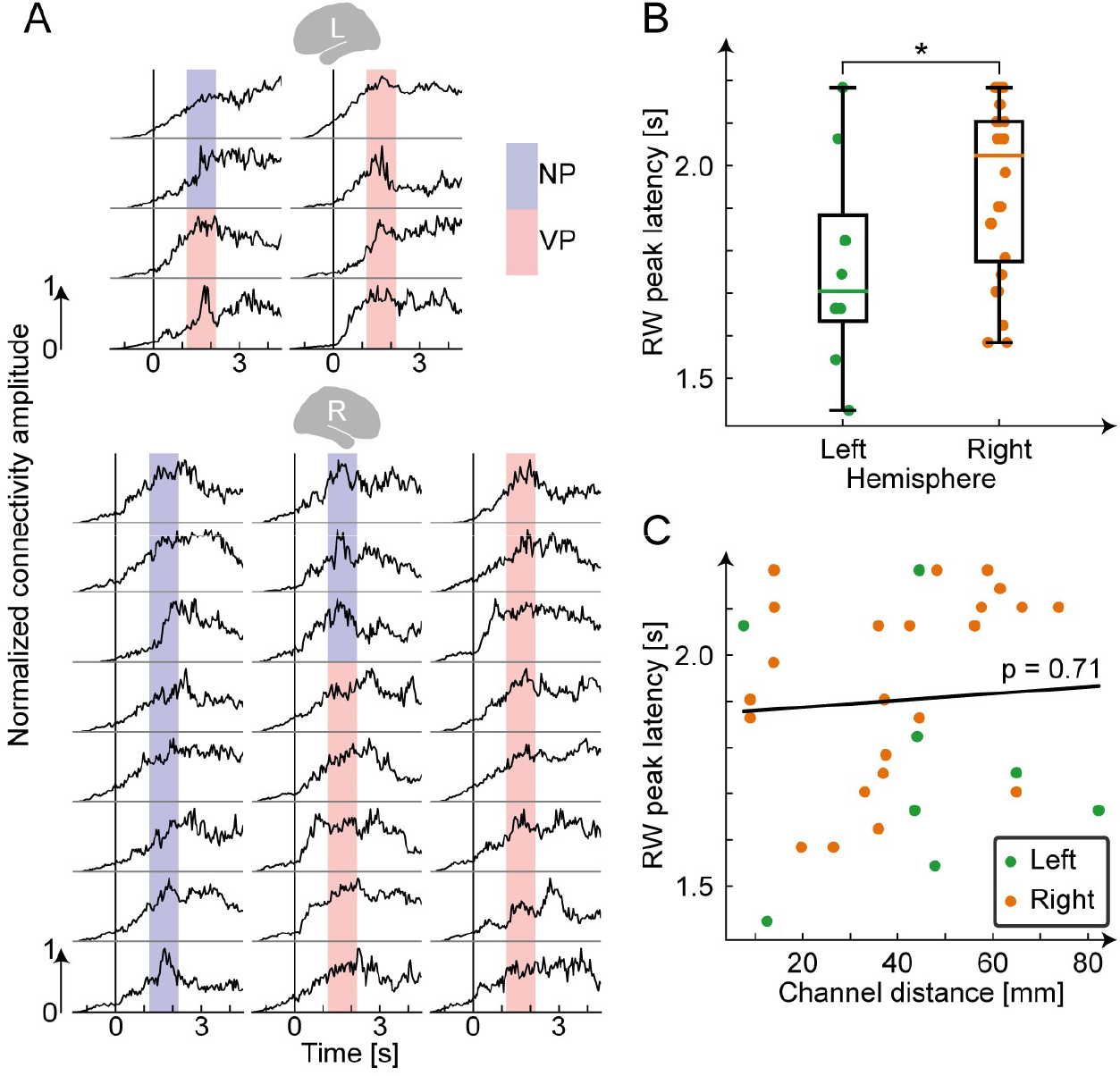
Syntax is faster in the dominant hemisphere. **(A)** Time series of the identified directed connections, in the dominant (L, top) and non-dominant (R, bottom) hemispheres. Each time series is normalized between 0 and 1. The 0 in the time axis is the start of the sentence, the coloured area represents the homophonous part (or response window - RW). Directed connections that are significant when listening Noun Phrases (NP) have this area coloured in blue, while those significant for Verb Phrases (VP) have this area highlighted in red. **(B)** Box plots of the latencies of the connectivity peaks during the RW in the dominant hemisphere (left) and in the non-dominant one (right). **(C)** Scatter plot of latencies of the peaks during the RW as function of the distances between channel pairs.

The peak latencies in the directed connections evoked by the homophonous syntagms did not correlate linearly with the distances between the recording contacts involved in those connections (Pearson’s *ρ* = 0.07, p = 0.71, **Figure 3C**). Moreover, distances between recording contacts implanted in the DH and NDH and participating in an active connection were not statistically different (Mann-Whitney U_8,24_ = 83, p = 0.29). Therefore, the difference in peak latencies was likely not due to the channel distribution in the two hemispheres, but rather solely to the syntactic processing time.

### Connectivity decodes homophonous phrases

The general neural connectivity estimated by the time-varying PDC was able to determine if the subject was waiting for the sentence (baseline), listening to the initial part of the sentence, to the homophonous phrase (RW), or its ending. We used a Long Short-Term Memory Network (LSTM) (Hochreiter & Schmidhuber, 1997) to classify the stimulus segments with single-trial accuracy equal to 83.75 % (**Figure 4A**).

**Figure 4.**
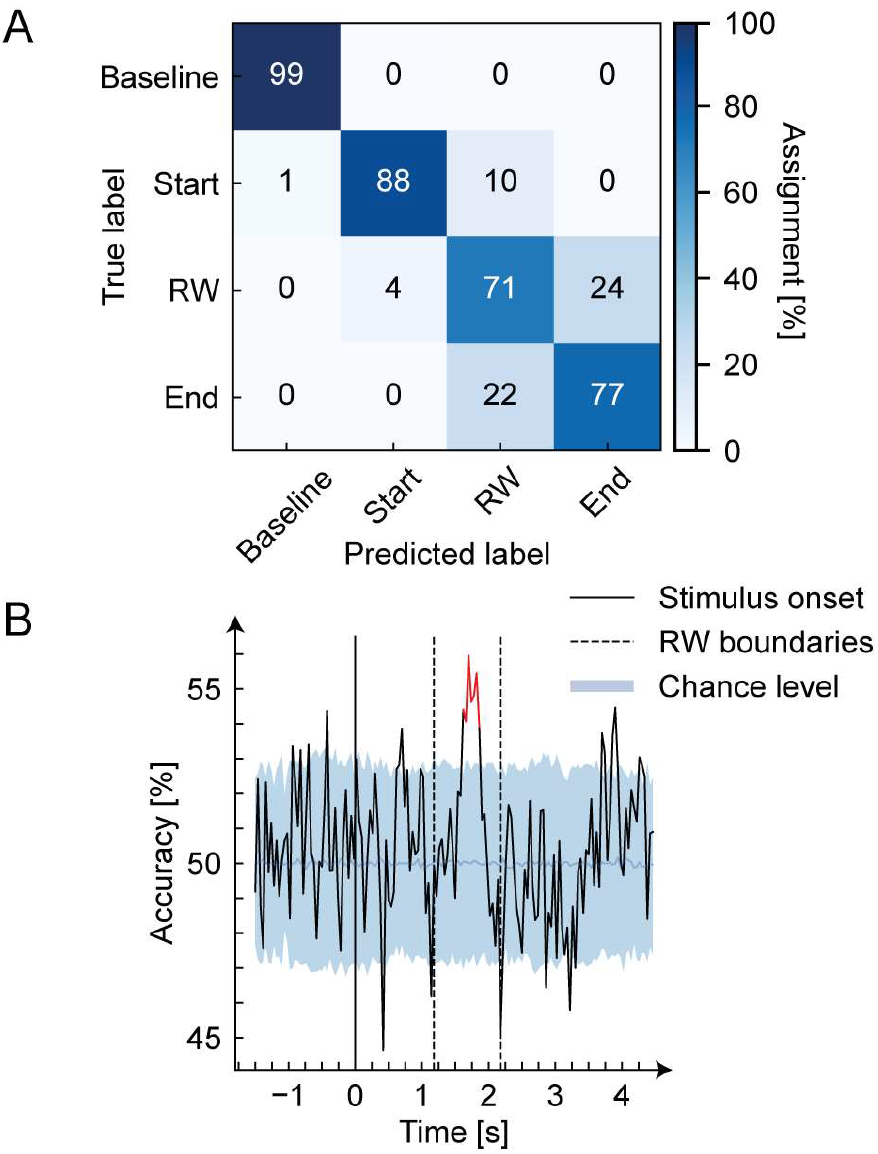
Connectivity decodes sentence structure. **(A)** Confusion matrix for the prediction of the stimulus phase. **(B)** Time-varying accuracy of the classification of noun phrases vs verb phrases. The blue line represents the median of the chance level, the boundaries of the light blue band are the 5th and 95th percentiles of the chance level distribution. The red part of the plot is the accuracy significantly above the chance level (p < 0.005).

We finally extracted time-dependent features only on the identified significant connections. We used a Support Vector Machine (SVM) (Cortes & Vapnik, 1995) to predict the syntactic content of the homophonous phrase in the sentence. The accuracy was significantly above chance during the RW phase (**Figure 4B**).

Both models were evaluated using a Leave-One-Subject-Out (LOSO) cross-validation.

## Discussion

Language comprehension and production, in particularly syntax processing, are complex and highly integrated tasks continuously carried out by our brain, seemingly without effort. Analysing their neural correlates thus requires sophisticated tools. One of the most promising techniques to identify the different neural processes underlying the syntactic operations leading to the processing of, for example, Noun Phrases or Verb Phrases is offered by directed connectivity evaluation related to the complexity of the large-scale networks. To our knowledge, this is the first time a difference in the connectivity elicited by NPs or VPs processing was identified.

Traditionally, the problem of understanding the neural correlates of syntax is approached by studying the effects of brain lesions or with syntax-related experimental tasks adminstered during neurophysiological and neuroimaging acquisitions contaminated by confounding factors such as phonology or semantics (Friederici et al., 2017; Vigliocco et al., 2011). Our approach is to leverage NP/VP homophonous phrases. The advantage of our solution is that we can factor out confounding factors by analyzing these homophonous phrases.

The shift from the analysis of isolated lexical elements such as bare Vs and Ns vs. syntactic units, namely VPs and NPs, is obviously a necessary step toward the goal of capturing syntactic information. Lexical elements in isolation contain linguistic information but these pieces of information are artificially expressed in single words whereas natural linguistic expressions always involve syntactic computation. In fact, the stimuli involved syntax in two directions: first, each homophonous phrase was syntactically connected with other words expressing a full-fledged sentence; second, each homophonous phrase contained very different syntactic structures. More specifically: in NPs the surfacing order of the two words composing them, namely an article and a noun, was the same as the underling structure composing it; in VPs, the situation is completely different and definitely more complex. In all VPs considered here a transformation called *cliticization* takes place. The order of the elements constituting it (a pronoun, playing the role of the object, and a verb) is reversed with respect to the canonical order in an SVO language like Italian; the canonical position of the object is to the right of the verb (Moro, 2016). All in all, the shift from V/N to VP/NP constitutes a necessary and relevant step towards the final goal of cracking the underlying code of human syntax.

The information carried by all the directed connections was able to discriminate between parts of the sentence. The syntactic category of the stimulus was discriminable only by looking at the significant connections, showing the importance of restricting the topology analysis on the few significant connections.

In recent years, there have been important technological and methodological advancements in perceived and imagined speech decoding (Martin et al., 2018; Panachakel & Ramakrishnan, 2021). Recent works focus on the classification of vowels (M. S. Mahmud et al., 2020; N. T. Duc & B. Lee, 2020), syllables (Archila-Meléndez et al., 2018; Brandmeyer et al., 2013; Correia et al., 2015), words (Ossmy et al., 2015; Proix et al., 2022; Vorontsova et al., 2021) and complete sentences (Chakrabarti et al., 2015; Zhang et al., 2012), distinguishing stimuli mainly at the semantic level. The most advanced online decoding techniques rely heavily on the articulatory representation of syllables and words in the motor and supplementary motor cortices (Anumanchipalli et al., 2019). However, this approach can only be applied to patients with intact motor commands, which represent a minority of the patients with speech impairment (Guenther et al., 2009; Wilson et al., 2020). Thus, other decoding strategies that rely on the brain regions that encode speech are needed (Proix et al., 2022).

Here, we decoded the acoustic stimuli exploiting 29 different speech-encoding cortical areas spanning the entire brain. Only recently such strategy has been used in the decoding of groups of syllables and words (Proix et al., 2022).

However, our approach relies on the time evolution of the connectivity values between recording contacts. This solution has the advantage of assuring high inter-subject generalizability as shown by the LOSO validation results: the connectivity features are independent of the location of the implanted leads, which may differ from subject to subject. Also, our method is well suited to be implemented in an online decoder. Moreover, the signals that drive the decoding are directly entangled to the syntactic representation of the stimuli rather than their phonological - and articular - components.

We believe that a decoding strategy that relies on multiple language-encoding cortical areas will drastically improve the performance of speech prostheses and may be the key missing piece for the development of this technology.

We showed that VPs processing, compared to NPs processing, elicited a significantly higher number of directed connections, linked together more brain structures both in the DH and in the NDH, and involved the activation of a wider cortical network. VPs processing was distributed beyond temporal lobes, pushing the information from sources located in the right frontal lobe and left insula, to sinks in both frontal lobes, anterior cingulate regions, and right insula. This suggests a greater network small-worldness for NPs, with a preference for short-range connections over long range ones.

Most of the literature converges on a more extended cerebral involvement in verb processing than for nouns (Lukic et al., 2021; Vigliocco et al., 2011). However, again, most evidence came from tasks requiring the processing of N/V as words in isolation: this is the first time an approach based on homophonous phrases, hence syntax, is used.

Temporal lobes (both in the DH and in the NDH) seem to be the main hub in which the syntactic operations leading to NPs or VPs are analyzed and processed. For NPs all the information flow started from these areas, while for VPs 3 out of 20 sources were placed outside the temporal lobes (with the one in the right occipital cortex very close to temporal areas). Also, sinks were mostly located in the temporal lobes. The important role of the temporal lobes, in particular of left posterior regions, in syntactic processing is supported by lesion and imaging evidence (Friederici et al., 2017; Matchin & Hickok, 2020). The comparison of the estimated directed connections with the CCEPs arising between recording contacts showed a partial discrepancy. While the structural connectivity underlying CCEPs is well known (e.g., the Human Connectome Project) (Van Essen et al., 2012), the functional and effective connectivity are patterns of highly heterogeneous causal relationships that may reflect processes occurring during many different temporal time scales (Honey et al., 2009; Keller et al., 2014; Matsui et al., 2011; Shmuel & Leopold, 2008; Vincent et al., 2007). The event-related causality identified here, is thus the expression of more complex neural processes, for which there are no unique a priori hypotheses.

Interestingly, recording contacts involved in a significant connection and showing at the same time CCEPs were implanted closed together than the pairs of channels without relevant CCEPs. Indeed, CCEPs may terminate their propagation early (Keller et al., 2014; Logothetis et al., 2010), which is in agreement with the description of CCEPs as supported by short-range local relations arising from direct hardwired connections via cortico-cortical or cortico-subcortico-cortical pathways (Matsumoto et al., 2004). This suggests that syntax-related processing relies mostly on long-range connections between cortical areas, expressing network-level neural synchronization supported by long-range, indirect structural pathways, typical of high-level cognitive processing (Salmelin & Kujala, 2006).

Earlier peaks in the connectivity time-series in the DH revealed that the syntax processing elicited by our stimuli started first in the temporal lobes of the left hemispheres and then spread to the right cortices. The directed links from DH to NDH that are necessary to transfer the information from one hemisphere to the other were not deemed significant because they were probably active during all sentence processing, and so they were masked during the search for the causal connections with the highest amplitude increase during the homophonous part of the stimulus. Also, only 1 subject out of 10 was explored in both hemispheres.

Focal lesion, behavioral, fMRI and electrophysiological studies provide converging evidence for a dominant role of one hemisphere (the left in right-handers and in the majority of left-handers) for most aspects of language processing (Tzourio-Mazoyer et al., 2017). Here we detected more significant connections arising in the NDH than in the DH. Previous studies claim that speech perception is the only aspect of bi-hemispheric language processing, even if the successive linguistic elaboration is carried mainly by the DH (Hickok & Poeppel, 2000; Poeppel et al., 2008). We hypothesize that the lesser number of significant connections in the DH may be due to the spread of the syntactic-related information to the NDH after it is first processed in the DH. However, future work is needed to better characterize the role of the NDH in syntax processing.

In conclusion, these results represent an important step forward in human language comprehension, contributing to the full characterization of syntactic processing. We showed a specific brain activity encoding a syntactic distinction, which is faster in the DH. Since, even from a purely formal point of view, syntactic processing cannot be compared with other computational systems, language-related or not (Chomsky, 2014; Moro, 2014b, 2014a), it is reasonable to conclude that the network highlighted here is not only specific but arguably it is uniquely dedicated to syntax. We prove that it is possible to decode the syntactic structure of a phrase by looking at the connections elicited by speech processing between multiple cortical areas. This could contribute to the future development of speech prostheses for speech impairment mitigation (Anumanchipalli et al., 2019).

## Acknowledgments

This work was financed by the Italian Ministry of Education, University and Research (MIUR) through PRIN-2017 ‘INSPECT’ (Project 2017JPMW4F) and by the Bertarelli Foundation.

## Author contributions

Conceptualization: S.F.C., A.M., and S.M.; Methodology: F.B., C.R., G.L.R., S.F.C., A.M., and S.M.; Software: A.C. and F.A.; Validation: A.C., F.A., P.d.O., and M.R.; Formal Analysis: A.C.; Investigation: P.d.O. and M.R.; Resources: G.L.R. and S.M.; Data Curation: A.C., F.A., P.d.O., and M.R.; Writing, Review and Editing: A.C., F.A., S.F.C., A.M., and S.M.; Visualization: A.C.; Supervision: G.L.R., S.F.C., A.M., and S.M.; Project Administration: S.F.C., A.M., and S.M.; Funding Acquisition: S.M.

## STAR Methods

### Resource availability

#### Lead Contact

Further information and requests for resources should be directed to and will be fulfilled by the lead contact, Fiorenzo Artoni.

#### Materials availability

This study did not generate new unique materials.

#### Data and code availability

All the data used for the study are available from the authors upon reasonable request.

This study adapts built-in MATLAB functions, the EEGLAB toolbox, the BrainNet Viewer toolbox, and several publicly available *Python* packages to handle, analyze and plot data. Custom code is available from the authors upon request.

Any additional information required to re-analyze the data reported in this paper is available from the lead contact upon request.

### Experimental model and subject detail

#### Human subjects

In total, 23 patients were recruited. All of them underwent surgical implantations of intracerebral electrodes for refractory epilepsy (Cossu et al., 2015) in the “Claudio Munari” Epilepsy Surgery Center of Milan, Italy (Cossu et al., 2005; Munari et al., 1994). The strategy of implantation was defined purely based on clinical needs, to locate the epileptogenic zone.

All patients completed all experimental sessions. During the 24h before the experimental recording, no seizure occurred, no alterations in the sleep/wake cycle were observed, and no additional pharmacological treatments were applied. No language or neuropsychological deficits were found in any patients. Also, no anatomical alterations were made evident by magnetic resonance. High-frequency stimulation (50 Hz, 3 mA, 5 sec) through SEEG electrodes was used to assess language dominance in all subjects. Two patients also underwent an fMRI study during a language task before the implantation of the electrodes.

Thirteen patients were excluded from the analysis. Eight of them exhibited pathological SEEG contacts. The others five patients showed no explored recording contacts with a task-related significant activation in our previous study (Artoni et al., 2020). Full demographic data are shown in **Table S4**.

A total of 2186 recording contacts (median 210, range 168-272) were implanted, divided into 164 electrodes (median 16.5, range 13-19). The number of contacts in the grey matter was 1439 (65.8%); 586 recording contacts in the language dominant hemisphere (DH). The DH was explored in 5 subjects (median electrodes 16, range 3-18; median contacts 210, range 25-225). The non-dominant hemisphere (NDH) was explored in 6 subjects (median electrodes 15, range 14-19; median contacts 208, range 182-272). SEEG exploration involved both hemispheres with a preference for the non-dominant side in 1 patient.

Overall, 68 electrodes were implanted in the temporal lobe (26 in DH, 42 in NDH), 43 in the frontal lobe (22 in DH, 21 in NDH), 22 in the central lobe (9 in DH, 13 in NDH), and 30 in the parieto-occipital region (9 in DH and 21 in NDH).

The present study received the approval of the Ethics Committee of ASST Grande Ospedale Metropolitano Niguarda (ID 939-2.12.2013) and informed consent was obtained from all participants.

### Methods details

#### Stimuli

The set of stimuli is based on three characteristics of Italian. First, some definite articles are pronounced exactly like some object clitic pronouns (such as [la] written as *la*; it can be both “the - fem.sing.” or “her - fem.sing.”). Second, the syntax of articles and clitic pronouns is very different: articles precede nouns, complements follow verbs, but object clitics are placed before the verb. Third, the Italian lexicon contains several homophonous pairs of nouns and verbs, such as [‘pɔrta] (written *porta*), which can either mean “door” or “brings”. A set of pairs of words such as [la ‘pɔrta] (written as *la porta*) can thus be interpreted either as a noun phrase (“the door”) or a verb phrase (“brings her”) depending on the syntactic context (homophonous phrases). For example, in PULISCE LA PORTA CON L’ACQUA (s/he cleans the door with water), *la porta* is a Noun Phrase (NP), while in DOMANI LA PORTA A CASA (tomorrow s/he brings her home), *la porta* is a Verb Phrase (VP).

To be sure to eliminate phonological and prosodical factors, the pronunciation of one homophonous phrase was copied in the syntactic counterpart. No other semantic or lexical distinction differentiated the two types of phrases.

The acoustic stimuli were recorded using a Sennheiser Microphone MH40P48, connected via a Firewire 400 to an Apple OSX 10.5.8 with a Motu Ultralight Mk3 sound card. The stimuli were edited and mastered using Audiodesk 3.02 and Peak Pro7, respectively. Files were generated in 16 bits, with a sampling frequency equal to 44.1 kHz; intensity was normalized to 0 Db and rendered in .wav format. All sentences were read by the same person, an Italian native speaker, male, 53 years old.

#### Surgical procedure and recording equipment

SEEG electrodes have a diameter of 0.8 mm. They contain 5 to 18 recording contacts, which are 2 mm long and spaced by 1.5 mm. The strategy of implantation was planned on 3D multimodal imaging and the electrodes were stereotactically implanted with robotic assistance. The position of every recording contact was assessed by registering a post-implantation Cone-Beam-CT (O-arm scanner, Medtronic, Minneapolis, Minnesota) to pre-implantation T1 weighted MR images.

SEEG sampling rate during the experiment was set to 1 kHz (patients 1-12) or 2 kHz (patients 13-23). Recordings were carried out using a 192-channels EEG-1200 (Neurofax, Nihon Kohden). All recording contacts were re-referenced to two leads in the white matter, in which electrical stimulations did not produce any manifestation.

#### Recording protocol

Each subject rested in a comfortable armchair. Stimuli were delivered using the software Presentation (Neurobehavioral Systems). Phrases were delivered via audio amplifiers at the minimum volume for words to be perceived with ease, according to the subject. During stimuli delivery, subjects gazed at a 27 inches cross on a screen. A synchronization TTL trigger spike was sent to the SEEG trigger port at the beginning of the sentence. Jitter and delays were lower than 1 ms. The experiment lasted around 30 minutes. At the end of each task, subjects were always able to correctly answer short questions about the stimuli. A camera was used to control for eye movement, silence, and any unexpected behavior from the patients.

#### Data pre-processing

An anti-aliasing band-pass filter (0.015-500 Hz) was applied at the hardware level. Recordings acquired at 2 kHz were down-sampled to 1 kHz. Artifacts and pathological interictal activity were controlled and removed by clinicians and scientists by visual inspection. Recordings were annotated with the events triggered by the beginning of each word in all stimulus sentences. Epochs were extracted from −1.5 s to 4.5 s time-locked to the beginning of each stimulus. The length of the epochs always ensured the inclusion of the complete stimulus presentation. Epochs with notable artifacts were rejected. Contacts in white matter were excluded from the subsequent analysis.

#### Cortico-cortical evoked potentials

During the presurgical evaluation, an effective connectivity of the explored brain areas was assessed for each subject by evaluating the Cortico-Cortical Evoked Potentials (CCEP) elicited by Single-Pulse Electrical Stimulation (SPES) (Matsumoto et al., 2017; Russo et al., 2021; Trebaul et al., 2018).

In the condition of eyes open resting wakefulness, SPES was delivered through each pair of adjacent contacts, with at 5 mA current intensity, a single pulse of 0.5 ms (biphasic rectangular stimuli of alternating polarity), at 1 Hz frequency, for 15 s.

The presence of CCEPs response following a SPES was visually verified by trained neurophysiologists.

#### Stimulus-evoked causality estimation

To estimate the stimulus-evoked directed connections, recording contacts were first divided into mini-regions of interest (mini-ROIs). Then, the Partial Directed Coherence (PDC), a measure deriving from the Granger causality framework (Baccalá & Sameshima, 2001; Geweke, 1982; Granger, 1969) was computed. Finally, a non-parametric statistical test was used to evaluate the significant connections elicited in the response window (RW), i.e., the part of interest of the stimulus (NP or VP). This stimulus-evoked causality estimation pipeline, designed for SEEG data, is proposed in (Cometa et al., 2021).

##### Mini ROI extraction

Recording contacts showing high correlation coefficients between their time series were combined into mini-ROIs. Specifically, mini-ROIs are groups of leads having an averaged across trials coefficient of determination *R^2^* > 0.8. The prototypical channel of a mini-ROI was selected as the one showing the highest linear correlation with the mini-ROI mean time series. Mini-ROIs grouping was performed independently for each subject. Most mini-ROIs were populated by just one channel, with the most numerous ones not being populated by more than 3 recording contacts. All leads contained in a single mini-ROI were spatially very close and always belonged to the same electrode.

##### Causality estimation

Within the Granger causality framework, a time series *x_j_*(*t*) causes another time series *x_i_*(*t*) if knowledge of past samples of *x_j_*(*t*) reduces the prediction error for the current sample of *x_i_*(*t*). The relation between *x_j_*(*t*) and *x_i_*(*t*) can be estimated by fitting a time-varying multivariate autoregressive (MVAR) model on ***X***(*t*):

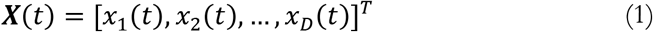

where *D* is the total number of channels.

The MVAR model assumes a linear relationship between the channels in ***X***(*t*) of the form:

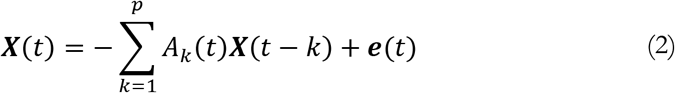

Where *A_k_*(*t*) is the time-varying *DxD* MVAR coefficients matrix, ***e***(*t*) is a white noise process with covariance matrix *W* and *p* is the model order. The *A_k_*(*t*) matrices were derived by using a general linear Kalman Filter (GLKF) (Milde et al., 2010). To estimate the model order *p*, the Bayesian information criterion (BIC) was used (Schwarz, 1978), resulting in *p* = 4 for all subjects.

After estimating, trial by trial, the *A_k_*(*t*) matrices, the single-trial time-varying *PDC*(*f*, *t*) (Astolfi et al., 2008) was computed.

To lower the computational complexity of the pipeline, PDC time samples were down-sampled by a factor of 40 (from 6000 samples to 150). Frequencies were averaged into overlapping frequency bins (width = 50 Hz, overlap = 25 Hz, range = 0 – 300 Hz).

Subsequent analysis was done only in the ultra-high gamma frequency range (150 – 300 Hz), i.e., on frequency bins from [125-175 Hz] to [250-300 Hz].

##### Significance during the homophonous phrase

All the next steps of the algorithm were independently applied for each syntactic structure (NPs or VPs), each subject, and each frequency band *f*. Linear interpolation time-warping was used to align the RW across all trials (Artoni et al., 2017; Do et al., 2021; Gwin et al., 2011; Nordin et al., 2019). Baseline correction was then carried out by dividing *PDC_ij_*(*f*, *t*), trial by trial and for each *i*, *j* (*i* ≠ *j*) couple independently, by its mean baseline value. The 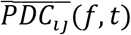 matrices were obtained by averaging *PDC_ij_*(*f*, *t*) over trials. The mean values of the 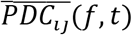 during the RW were calculated for each pair *i*, *j* (*i* ≠ *j*) of channels. These mean values were compared against a null distribution: to generate the null (permutation) distribution and to control for false discovery rate (Maris & Oostenveld, 2007; Nichols & Holmes, 2002) the time samples of the 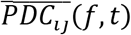 were shuffled 1000 times and the mean values during the RW were re-computed for each permutation. The maximum mean value across all channel couples was retained for each permutation.

A *p-value* for each recording contact pair was calculated by comparing the original mean connectivity value in the RW with the permutation distribution. The *p-value* was the number of instances in the null distribution that were greater than the mean RW causality. Significance was then assigned to connections between pairs of leads whose *p-value* was below a certain threshold. The threshold was set to 0.33, being the lowest one that allowed the arising of at least one significant connection for either NPs or VPs in every subject, in at least one of the considered frequency bins.

#### Latency analysis

To detect the peaks in connectivity during the RW of the stimuli, the average connectivity time series were first smoothed. A Savgol filter was used (Guiñón et al., 2007). The polynomial order was set to 2, with 9-samples long windows. The window size was chosen as the knee of the curve formed by the sum of absolute differences between the smoothed time series and the raw ones for different window lengths. The latencies were defined as the time instant at which the maximum of each smoothed time series occurred, within the homophonous phrase interval.

#### Cortical surface plotting

Mini-ROIs (**Figure 1B**), active directed connections (**Figure 2A**), and active cortical areas (**Figure 4A**) were graphically represented using the *BrainNet Viewer* toolbox for Matlab (Xia et al., 2013). Plotting was done using MNI coordinates on a FreeSurfer *fsaverage* template (Fischl, 2012; Wu et al., 2018).

#### Decoding

##### Response window prediction

The prediction of the phase of the stimulus was carried out on a trial-by-trial basis. All the connections were used. The time-varying connectivity amplitudes were divided into overlapping bins of size 20 samples and step 1 sample. These were fed to a Long Short-Term Memory network (LSTM) (Hochreiter & Schmidhuber, 1997) together with the labels corresponding to the stimulus phase (baseline, sentence start, RW, sentence ending) of the last sample of the overlapping windows.

The training was carried out using a Leave-One-Subject-Out (LOSO) cross-validation procedure. For each fold of the LOSO cross-validation, 2 trials of the training set were removed and used as the validation set. The decoder hyperparameters were optimized according to the performance on the validation set.

A weighted version of the categorical cross-entropy (Martín Abadi et al., 2015; Y. Ho & S. Wookey, 2020) was used as the loss function to minimize during the training of the LSTM, with the weights for each class inversely proportional to the length of the stimulus phase.

The accuracy was obtained by averaging the accuracies across all folds of the LOSO cross-validation. Code implementation was based on the *TensorFlow* package for python (Martín Abadi et al., 2015).

##### Syntactic content decoding

The prediction of the content of the homophonous phrases (NP vs VP) was carried out on a trial-by-trial basis. Only the significant connections were selected, regardless of whether the connections were significant during NPs or VPs processing. For each time point, a number of values equal to the number of significant connections were thus retained, corresponding to the amplitudes of the significant connections during that instant. A total of 7 features were then calculated for each time point: the statistical moments up to order 4, the median, the maximum, and the range (the difference between the maximum and the minimum).

A Support Vector Machine (Cortes & Vapnik, 1995) with a radial basis function kernel was trained for each time point. The training was carried out using a nested cross-validation procedure: (i) LOSO cross-validation was used to split the dataset into training and test set, and (ii) for each fold of the LOSO cross-validation, 10 fold cross-validation was used to furtherly divide the training set into training and validation set.

The inner validation loop was used to optimize the decoder hyperparameters and to perform feature selection through the minimum redundancy maximum relevance (Radovic et al., 2017) algorithm.

The time-varying accuracy was obtained by averaging the accuracies across all folds of the LOSO cross-validation procedure.

For each time point, the predicted labels were compared 1000 times with 1000 shuffled versions of the test set labels (NP or VP) to calculate the chance level. The procedure was repeated for each fold of the LOSO cross-validation, resulting in a null distribution of 1000 x (number of fold) accuracy values. An exact p-value was obtained by comparing the original accuracy with the null distribution.

The time-varying p-values were corrected for the multiple comparisons using a cluster-size-based statistical non-parametric mapping approach (Nichols & Holmes, 2002) and deemed significant if lower than *α* = 0.05.

Code implementation was based on the *scikit-learn* package for python (Pedregosa et al., 2011).

### Quantification and statistical analysis

The non-normality of the data undergoing statistical testing was assessed using Shapiro-Wilk tests (Shapiro & Wilk, 1965). Sizes *n1* and *n2* of the independent samples undergoing Mann-Whitney tests (Neuhäuser, 2011) and the associated *U* statistics are reported in the Results Section as *U_n1,n2_* = *U*. Statistical significance level α was 0.05. The inter-hemispheric significant connection that arose in one subject was not considered in the tests comparing connections in the DH versus connections in the NDH. Tests were computed using the *scipy* package for *Python* (Virtanen et al., 2020).

## KEY RESOURCES TABLE

| REAGENT or RESOURCE | SOURCE | IDENTIFIER |
| --- | --- | --- |
| Software and Algorithms |  |  |
| MATLAB | MathWorks | https://www.mathworks.com/ |
| EEGLAB toolbox for MATLAB | Swartz Center for Computational Neuroscience | [EEGLAB (ucsd.edu)](http://EEGLAB.ucsd.edu) |
| Python | Python | [Welcome to Python.org](http://Welcome.to.Python.org) |
| Scipy ecosystem for Python | Scipy | SciPy.org |
| scikit-learn | Scikit-Learn | scikit-learn.org |
| Pytorch | Google AI | [10.5281/zenodo.4724125](https://doi.org/10.5281/zenodo.4724125) |
| BrainNet Viewer toolbox for MATLAB | Xia et al., 2013 | [NITRC: BrainNet Viewer: Tool/Resource Info](http://NITRC.org/BrainNet-Viewer/Tool/Resource-Info) |
| Presentation | Neurobehavioral Systems | [Neurobehavioral Systems (neurobs.com)](http://Neurobehavioral-Systems.neurobs.com) |

