## Supplemental Tables for "Event-related causality in Stereo-EEG discriminates syntactic processing of noun phrases and verb phrases"

| Subj. ID | From | To | CCEP |
|---|---|---|---|
| **NOUN Phrases** | | | |
| s03 | R parahippocampal gyrus (39, -21, -30) | R planum polare (44, -23, 6) | N |
| s03 | R superior temporal gyrus (58, -24, 5) | R postcentral gyrus (inferior third) (46, -10, 14) | Y |
| s09 | R superior temporal gyrus (62, -12, 1) | R transverse temporal gyrus (Heschl) (41, -24, 10) | Y |
| s09 | R temporal pole (25, 10, -35) | R transverse temporal gyrus (Heschl) (45, -26, 10) | N |
| s12 | R inferior temporal sulcus (59, -24, -15) | R Inferior frontal gyrus (pars triangularis) (56, 26, 12) | N |
| s12 | R hippocampus (26, -18, -16) | R anterior cingulate gyrus (9, 32, 22) | N |
| s12 | R temporal pole (22, 2, -31) | R inferior frontal gyrus (pars orbitalis) (43, 43, -16) | N |
| s13 | L hippocampus (-31, -31, -9) | L hippocampus (-24, -31, -9) | Y |
| s13 | L planum polare (-47, 16, -16) | L transverse temporal gyrus (Heschl) (-38, -20, 9) | N |
| **VERB Phrases** | | | |
| s03 | R superior temporal gyrus (54, 11, -11) | R inferior temporal gyrus (63, -22, -27) | Y |
| s06 | L superior circular sulcus of insula (-36, 6, 6) | L superior frontal gyrus (-15, 67, 15) | Y |
| s09 | R inferior temporal gyrus (56, -21, -29) | R transverse temporal gyrus (Heschl) (41, -24, 10) | N |
| s10 | L middle temporal gyrus (-55, -63, 7) | L posterior cingulate gyrus (-8, -52, 8) | N |
| s10 | L superior temporal gyrus (-57, -16, 4) | L planum polare (-45, -17, 2) | Y |
| s11 | L middle temporal gyrus (-64, -16, -16) | L suborbital sulcus (-4, 41, -13) | N |
| s12 | R hippocampus (26, -18, -16) | R anterior cingulate gyrus (10, 52, 3) | N |
| s12 | R precentral sulcus (inferior port) (59, 6, 17) | R superior frontal gyrus (12, 52, 22) | Y |

| | | | |
|---|---|---|---|
| s12 | R temporal pole<br>(22, 2, -31) | R anterior cingulate gyrus<br>(10, 52, -1) | N |
| s13 | L planum polare<br>(-42, 16, -15) | L transverse temporal gyrus (Heschl)<br>(-38, -20, 9) | N |
| s13 | L planum polare<br>(-47, 16, -16) | L transverse temporal gyrus (Heschl)<br>(-42, -21, 9) | N |
| s17 | R superior temporal sulcus<br>(58, -27, -11) | R posterior portion of lateral fissure<br>(32, -24, 10) | N |
| s17 | R middle temporal gyrus<br>(65, -14, -18) | R posterior portion of lateral fissure<br>(32, -24, 10) | N |
| s21 | R middle temporal gyrus<br>(67, -27, -8) | R long gyri of insula<br>(44, 2, -8) | N |
| s21 | R lingual gyrus<br>(19, -59, -6) | R middle temporal gyrus<br>(67, -27, -8) | N |
| s23 | R superior temporal gyrus<br>(59, -2, -5) | L transverse temporal gyrus (Heschl)<br>(-49, -13, 4) | N |
| **NOUN and VERB Phrases** | | | |
| s03 | R superior temporal gyrus<br>(58, -24, 5) | R planum polare<br>(44, -23, 6) | Y |
| s17 | R middle temporal gyrus<br>(65, -27, -12) | R posterior portion of lateral fissure<br>(37, -24, 10) | N |
| s21 | R transverse temporal gyrus (Heschl)<br>(49, -22, 3) | R circular sulcus of insula<br>(40, -21, 3) | Y |
| s21 | R transverse temporal gyrus (Heschl)<br>(49, -22, 3) | R superior temporal sulcus<br>(53, -29, -8) | N |

**Table S1. Directed connections.** List of the significant directed connections in the whole population of patients. Subject ID, presence or absence of CCEP, brain region and MNI coordinates of the source, brain region and MNI coordinates of the sink are indicated for each connection. Subj: Subject. R: Right. L: Left. CCEP: Cortico-cortical Evoked Potential. Y: Yes. N: No.

| Subj. IDs | Source – Brain Region |
|---|---|
| **NOUN and VERB Phrases** | |
| s03, s17 | R middle temporal gyrus |
| s21 | R transverse temporal gyrus (Heschl) |
| s03, s09 | R superior temporal gyrus |
| s13 | L planum polare |
| s09, s12 | R temporal pole |
| s12 | R hippocampus |
| **NOUN Phrases** | |
| s03 | R parahippocampal gyrus |
| s12 | R inferior temporal sulcus |
| s13 | L hippocampus |
| **VERB Phrases** | |
| s06 | L superior circular sulcus of insula |
| s09 | R inferior temporal gyrus |
| s10, s11 | L middle temporal gyrus |
| s10 | L superior temporal gyrus |
| s12 | R precentral sulcus (inferior port) |
| s17 | R superior temporal sulcus |
| s21 | R lingual gyrus |

**Table S2. Sources.** List of brain zones (Destrieux atlas) acting as sources in the significant connections. Subj: Subjects. R: Right. L: Left.

| Subj. IDs | Sink - Brain Region |
|---|---|
| **NOUN and VERB Phrases** | |
| s21 | R circular sulcus of insula |
| s03 | R planum polare |
| s17 | R posterior portion of lateral fissure |
| s09 | R transverse temporal gyrus (Heschl) |
| s13, s23 | L transverse temporal gyrus (Heschl) |
| s12 | R anterior cingulate gyrus |
| **NOUN Phrases** | |
| s03 | R postcentral gyrus (inferior third) |
| s12 | R inferior frontal gyrus (pars triangularis) |
| s12 | R inferior frontal gyrus (pars orbitalis) |
| s13 | L hippocampus |
| **VERB Phrases** | |
| s21 | R middle temporal gyrus |
| s03 | R inferior temporal gyrus |
| s21 | R long gyri of insula |
| s12 | R superior frontal gyrus |
| s06 | L superior frontal gyrus |
| s21 | R superior temporal sulcus |
| s10 | L planum polare |
| s11 | L suborbital sulcus |
| s10 | L posterior cingulate gyrus |

**Table S3. Sinks.** List of brain zones (Destrieux atlas) acting as sinks in the significant connections. Subj: Subjects. R: Right. L: Left.

| | |
|---|---|
| Number of patients | 10 |
| Gender | 5 male and 5 female |
| Median age at recording | 32 (range 17-44) |
| Median years of scholarization | 13 (range 0-17) |
| Median age of onset of epilepsy | 16 (2-30) |
| Median duration | 15 (3-30) |

**Table S4. Demographic data.** Summary of demographic data of patients for whom connectivity was estimated. SEEG: Stereo-Electro-Encephalography.
